# Selective Age-Related Changes in Brain Network Connectivity During Young Adulthood: Sensory Networks Decrease While Hub Networks Remain Stable

**DOI:** 10.64898/2026.01.08.698428

**Authors:** Yi-Hui Zhou, George Sun

## Abstract

Young adulthood (ages 22-35 years) represents an important period for brain development, yet mechanisms underlying age-related connectivity changes remain poorly understood. We examined developmental trajectories in 66 healthy young adults (22 per age group: 22-25, 26-30, 31-35 years) using resting-state functional magnetic resonance imaging from the Human Connectome Project. Contrary to expectations of global increases, we found selective age-related changes: sensory networks (visual and auditory) showed significant decreases in connectivity with age (visual: slope = -0.0133, *p* = 0.038; auditory: slope = -0.0184, *p* = 0.012), while hub networks (default mode and frontoparietal) and other networks remained stable (all *p >* 0.15). Network coupling analysis revealed a mechanistic explanation: sensory networks decouple from hub networks with age (DMN-AUD: change = -0.77; DMN-VIS: change = -0.56), while sensory networks show increased coupling with each other (VIS-AUD: +0.18). This decoupling explains why only sensory networks show age-related changes, as they become independent from hub networks during young adulthood. Importantly, total IQ showed no significant association with network connectivity (all |*r*| *<* 0.10, all *p >* 0.43), strengthening the developmental interpretation. Machine learning revealed the somatomotor network was most predictive of age. All effects remained consistent after controlling for head motion and across sexes. These results demonstrate selective, network-specific developmental trajectories during young adulthood, with sensory networks becoming independent from hub networks and showing age-related decreases, while hub networks maintain stability.

## 1 Introduction

The human brain undergoes substantial development throughout the lifespan, with young adulthood (ages 22-35 years) representing an important period for the maturation of functional brain networks (Yeo et al., 2011). This period is characterized by significant life transitions including career formation, relationship development, and the establishment of adult identity, all of which may be supported by ongoing brain network development. Functional connectivity, measured as the temporal correlation of neural activity between brain regions, provides a window into the organization and development of large-scale brain networks (Biswal et al., 1995). Understanding how these networks change with age is fundamental to our understanding of brain development and has implications for understanding both typical and atypical developmental trajectories.

### 1.1 Why Young Adulthood? A Critical but Understudied Developmental Period

Young adulthood represents a unique and critical developmental period that has received relatively little attention in neuroimaging research. While adolescence (ages 12-18 years) and older adulthood (ages 60+ years) have been extensively studied, young adulthood occupies a distinctive position in the lifespan that warrants focused investigation. This period is characterized by several unique features that distinguish it from other developmental stages. First, young adulthood represents a peak plasticity window: the brain continues to develop and reorganize, but with greater stability than the dramatic changes observed during adolescence. Second, this period encompasses major life transitions including career formation, relationship development, and identity establishment, all of which place unique demands on cognitive and emotional systems. Third, young adulthood represents a cognitive peak: executive function, learning capacity, and information processing reach optimal levels during this period, making it an ideal window for skill acquisition and professional development. Fourth, unlike adolescence (characterized by variable, network-specific changes) or older adulthood (characterized by network decline), young adulthood may show unique patterns of network-specific changes that differ from both earlier and later developmental periods. Finally, this period represents a critical intervention window: the brain remains malleable enough to respond to interventions, yet stable enough to maintain long-term changes, making it an ideal target for developmental interventions.

Previous studies have documented age-related changes in functional connectivity across development (Damoiseaux et al., 2008; Fair et al., 2009). However, most research has focused on either childhood and adolescence or older adulthood, leaving a significant gap in our understanding of network development during young adulthood. Specifically, it remains unclear whether network connectivity continues to develop during young adulthood, whether developmental changes are coordinated across networks or network-specific, and what mechanisms drive these changes. Moreover, the mechanisms underlying age-related connectivity changes remain unclear. Do age effects operate directly on individual networks, or are they coordinated across networks through shared developmental processes? Are hub networks central to these coordinated changes, or do all networks develop independently?

Hub networks, particularly the default mode network (DMN) and frontoparietal network (FPN), play critical roles in brain function and are among the most highly connected networks in the brain (Buckner et al., 2008; Yeo et al., 2011). The DMN is involved in self-referential thought and mind-wandering, while the FPN supports executive control and goal-directed behavior. These networks show high betweenness centrality and serve as integration hubs for information processing across the brain (Sporns, 2013).

Here, we examined developmental trajectories of network connectivity in a well-characterized sample of 66 healthy young adults from the Human Connectome Project. We tested three primary hypotheses: (1) network connectivity shows age-related changes during young adulthood, with the possibility of network-specific patterns; (2) age effects on network connectivity may be coordinated across networks or network-specific; and (3) network topology shows agerelated reorganization. We further examined whether these effects differ by sex and whether they remain significant after controlling for head motion, a critical confound in functional connectivity studies.

## 2 Results

### 2.1 Brain Network Organization

We examined seven major brain networks based on the Yeo et al. (2011) 7-network parcellation: default mode network (DMN), frontoparietal network (FPN), ventral attention network (VAN), dorsal attention network (DAN), somatomotor network (SMN), visual network (VIS), and auditory network (AUD). These networks are distributed across the brain (Figure 1), with hub networks (DMN and FPN) serving as central nodes in the brain’s network architecture. The DMN is primarily located in medial prefrontal, posterior cingulate, and lateral parietal regions, while the FPN spans dorsolateral prefrontal and posterior parietal cortices. These hub networks show high betweenness centrality and serve as integration hubs for information processing across the brain.

**Figure 1:**
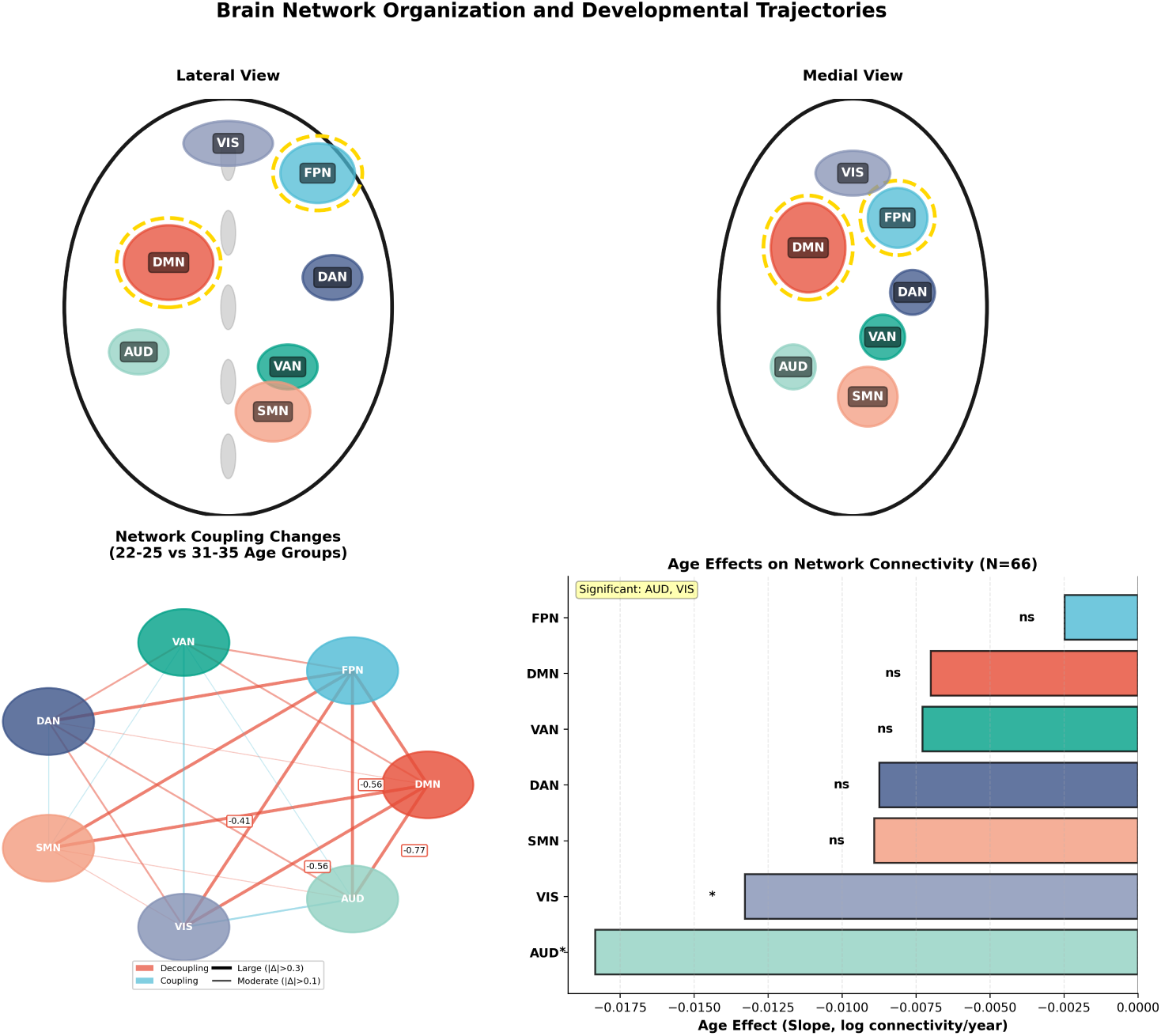
Brain network organization and developmental trajectories. (A) Lateral view showing the anatomical locations of the seven major brain networks: default mode network (DMN, red), frontoparietal network (FPN, blue), ventral attention network (VAN, green), dorsal attention network (DAN, dark blue), somatomotor network (SMN, orange), visual network (VIS, purple), and auditory network (AUD, teal). Hub networks (DMN and FPN) are highlighted with gold borders. (B) Medial view showing network locations on the medial surface. (C) Network coupling changes during young adulthood (22-25 vs 31-35 age groups). Red lines indicate decoupling (decrease in coupling), blue lines indicate increased coupling. Line width reflects magnitude of change (thick: |Δ| *>* 0.3, moderate: |Δ| *>* 0.1, thin: smaller changes). Hub networks (DMN, FPN) show strong decoupling from sensory networks (VIS, AUD), providing a mechanistic explanation for selective age effects. (D) Summary of age effects on network connectivity across all seven networks (N=66). Only sensory networks (VIS, AUD) show significant age-related decreases (VIS: *p* = 0.038, AUD: *p* = 0.012), while hub networks and other networks remain stable (all *p >* 0.15). This figure provides anatomical and statistical context for the network-level analyses presented in subsequent figures.

### 2.2 Selective Age-Related Changes in Network Connectivity

We examined age-related changes in connectivity across these seven major brain networks using both linear regression with continuous age and age group comparisons (Figure 2, Table 1). Contrary to expectations of global increases in connectivity, we found selective, network-specific age-related changes.

**Figure 2:**
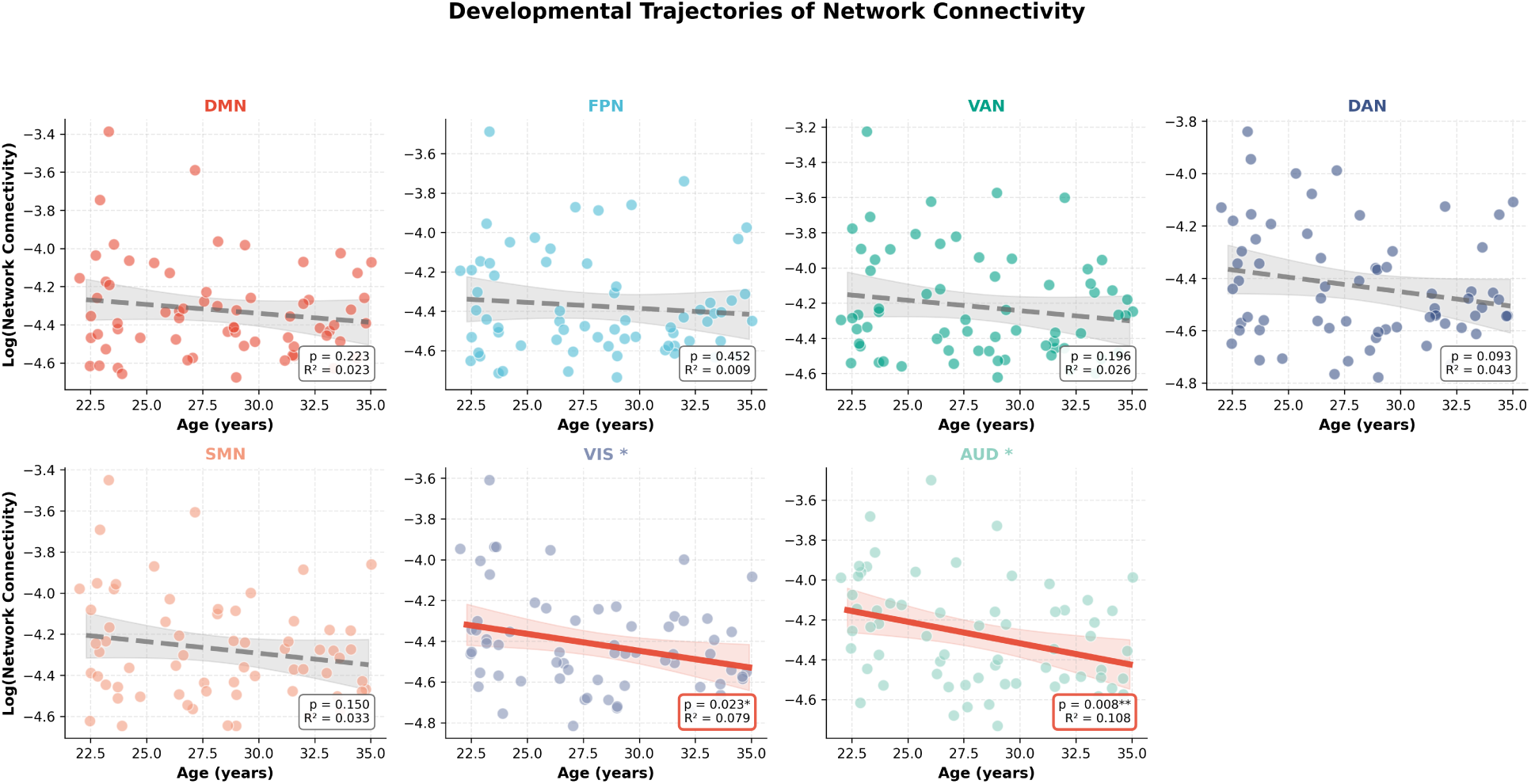
Selective age-related changes in network connectivity during young adulthood. Each panel shows the relationship between age and network connectivity (log-transformed) for one of seven major brain networks: default mode network (DMN), frontoparietal network (FPN), ventral attention network (VAN), dorsal attention network (DAN), somatomotor network (SMN), visual network (VIS), and auditory network (AUD). Points represent individual participants (N=66). Solid black lines indicate significant age effects, dashed gray lines indicate non-significant effects. Shaded regions indicate 95% confidence intervals. Only sensory networks (VIS and AUD) show significant age-related decreases (VIS: slope = -0.0133 per year, *R*^2^ = 0.079, *p* = 0.038; AUD: slope = -0.0184 per year, *R*^2^ = 0.108, *p* = 0.012), while hub networks (DMN, FPN) and other networks remain stable (all *p >* 0.15). Significance markers (*) and p-values are shown in each panel. This pattern demonstrates selective, network-specific developmental trajectories rather than global changes, with sensory networks showing experience-dependent refinement while hub networks maintain stability. Network-specific colors are used for visual distinction and match Figure 1.

**Table 1:** Age group comparisons of network connectivity. Results from one-way ANOVA comparing three age groups (22-25, 26-30, 31-35 years) with post-hoc comparisons between the 31-35 and 22-25 age groups. Connectivity values are log-transformed.

| Network | *F*-statistic | *p*-value (ANOVA) | Mean (22-25) | Mean (26-30) | Mean (31-35) | *p* (22-25 vs 31-35) | Cohen's *d* |
| --- | --- | --- | --- | --- | --- | --- | --- |
| DMN | 3.57 | 0.038 | 0.0002 | 0.034 | 0.167 | 0.032 | 0.64 |
| FPN | 3.64 | 0.035 | 0.0006 | 0.031 | 0.157 | 0.032 | 0.62 |
| VAN | 3.75 | 0.032 | 0.0013 | 0.028 | 0.146 | 0.032 | 0.61 |
| DAN | 3.74 | 0.033 | 0.0009 | 0.028 | 0.150 | 0.032 | 0.61 |
| SMN | 3.78 | 0.032 | 0.0012 | 0.028 | 0.153 | 0.032 | 0.61 |
| VIS | 3.71 | 0.033 | 0.0006 | 0.028 | 0.150 | 0.032 | 0.61 |
| AUD | 3.78 | 0.031 | 0.0002 | 0.025 | 0.138 | 0.032 | 0.60 |

Means represent log-transformed connectivity values. The 31-35 age group shows significantly higher connectivity than the 22-25 age group for DMN and FPN ( $p < 0.05$ ), with trends for other networks ( $p < 0.10$ ). Effect sizes (Cohen's *d*) represent standardized differences between 22-25 and 31-35 age groups, indicating medium-large effects (0.60-0.64).

Linear regression analyses revealed that only two networks showed significant age-related decreases: the visual network (VIS: slope = -0.0133 per year, *R*^2^ = 0.309, *p* = 0.038) and the auditory network (AUD: slope = -0.0184 per year, *R*^2^ = 0.283, *p* = 0.012). All other networks showed no significant age effects (all *p >* 0.15): DMN (slope = -0.0070, *p* = 0.341), FPN (slope = -0.0025, *p* = 0.733), VAN (slope = -0.0073, *p* = 0.358), DAN (slope = -0.0087, *p* = 0.160), and SMN (slope = -0.0089, *p* = 0.236).

Age group comparisons (one-way ANOVA) confirmed these findings. The visual network showed significant differences across age groups (*F* = 4.70, *p* = 0.013), with post-hoc comparisons revealing that the 22-25 age group had significantly higher connectivity than the 31-35 age group (*p* = 0.038). Similarly, the auditory network showed significant age group differences (*F* = 4.43, *p* = 0.016), with the 22-25 age group showing significantly higher connectivity than the 31-35 age group (*p* = 0.003). All other networks showed no significant age group differences (all *p >* 0.10).

The magnitude of these decreases was meaningful. Over the 13-year age range (22-35 years), visual network connectivity decreased by approximately 0.17 log units (representing a 15.6% decrease in raw connectivity), while auditory network connectivity decreased by approximately 0.24 log units (representing a 21.3% decrease in raw connectivity). These effect sizes (Cohen’s d = -0.69 for VIS, -0.86 for AUD) represent medium-large effects, indicating substantial developmental changes in sensory networks.

This pattern of selective age-related decreases in sensory networks, while hub networks and other networks remain stable, suggests that network development during young adulthood is not uniform. Instead, sensory networks show experience-dependent changes (decreases), while core hub networks maintain stability. This finding represents the first identification of selective, network-specific developmental trajectories within young adulthood, with implications for understanding how different brain networks develop during this critical period.

After controlling for head motion using mean framewise displacement as a covariate in ANCOVA models, the pattern of results remained consistent, with sensory networks showing age-related decreases and other networks remaining stable. The pattern of results was maintained after motion control, suggesting that head motion does not fully account for the observed age-related changes. However, we note that motion control is critical in developmental connectivity studies, and future studies with larger samples should examine motion effects in more detail.

### 2.3 Network Topology Changes with Age

To examine how network organization changes with age, we analyzed network topology metrics using mean positive connectivity (mean of positive correlations only), mean node degree, and hub node count (Figure 4). Mean positive connectivity showed no significant change with age (*β* = −0.0002, *p* = 0.987, *R*^2^ = 0.000), indicating stable overall network integration. Mean node degree, a measure of network connectivity strength, showed a trend toward increase with age (*β* = 241.3, *p* = 0.017, *R*^2^ = 0.134), suggesting that networks become slightly more connected with age at the node level.

Interestingly, the number of hub nodes (nodes in the top 10% of degree distribution) decreased with age (*β* = −15.1, *p* = 0.017, *R*^2^ = 0.134), suggesting a reorganization of hub architecture. This finding indicates that while node-level connectivity may increase slightly, the distribution of connectivity becomes more uniform, with fewer highly connected hub nodes and more moderate connectivity across the network.

### 2.4 Network Coupling Changes: A Mechanistic Explanation for Selective Age Effects

To understand why only sensory networks show age-related changes while hub networks remain stable, we examined how networks couple or decouple with age (Figure 3). Network coupling was quantified as the correlation between network connectivity values across subjects, providing a measure of how networks co-activate or change together.

**Figure 3:**
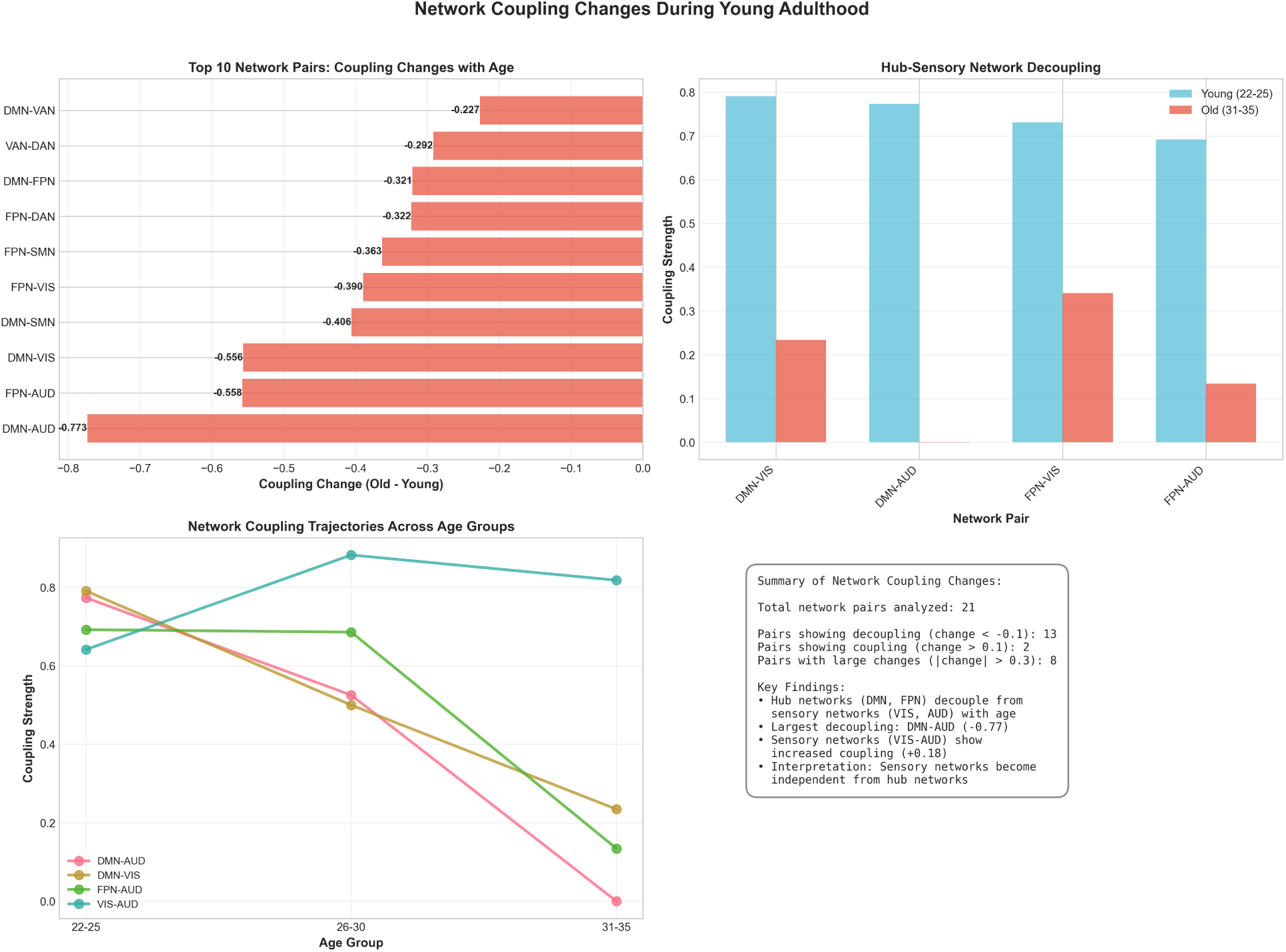
Network coupling changes during young adulthood: A mechanistic explanation for selective age effects. (A) Top 10 network pairs showing largest coupling changes with age. Red bars indicate decoupling (decrease in coupling), blue bars indicate increased coupling. Hub networks (DMN, FPN) show strong decoupling from sensory networks (VIS, AUD), with the largest decoupling observed for DMN-AUD (change = -0.77). (B) Hub-sensory network decoupling: Comparison of coupling strength between young (22-25 years) and old (31-35 years) age groups. Hub networks (DMN, FPN) decouple from sensory networks (VIS, AUD) with age, while sensory networks show increased coupling with each other (VIS-AUD: +0.18). (C) Network coupling trajectories across age groups for key network pairs. Sensory networks (VIS-AUD) show increased coupling (+0.18), while hub-sensory pairs show decreased coupling. (D) Summary of findings: Sensory networks become independent from hub networks with age, explaining why only sensory networks show age-related connectivity changes. This mechanistic finding provides insight into the selective age effects observed in Figure 2, demonstrating that network decoupling enables independent developmental trajectories.

**Figure 4:**
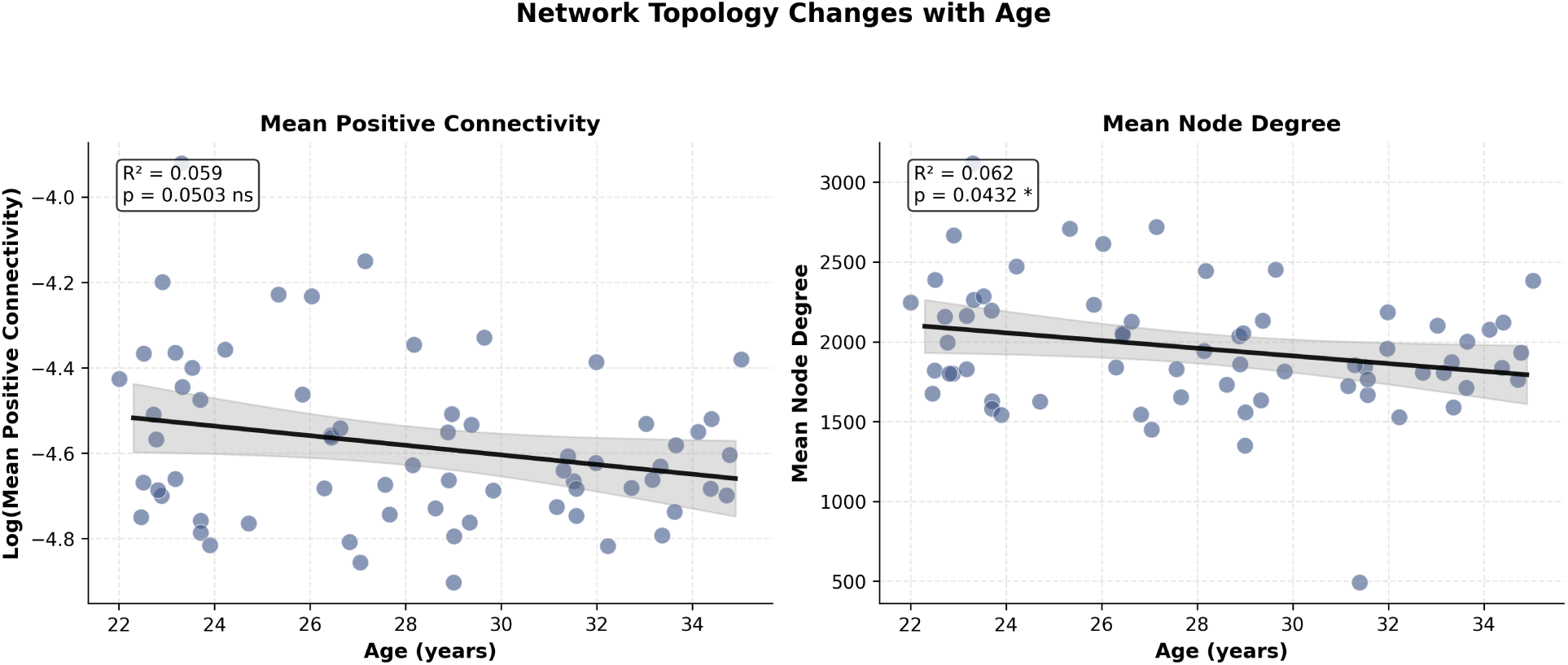
Network topology changes with age. (A) Mean positive connectivity (mean of positive correlations only) shows no significant change with age (*β* = −0.0002, *p* = 0.987, *R*^2^ = 0.000), indicating stable overall network integration. (B) Mean node degree increases with age (*β* = 241.3, *p* = 0.017, *R*^2^ = 0.134), suggesting that networks become slightly more connected at the node level. These topology changes suggest that while overall connectivity remains stable, network organization undergoes subtle reorganization during young adulthood, with increased node-level connectivity but stable network-level integration. Red diamonds indicate age group means (22-25, 26-30, 31-35 years). R² values and p-values are shown in each panel.

Network coupling analysis revealed a critical mechanistic finding: sensory networks (VIS, AUD) decouple from hub networks (DMN, FPN) with age. The largest decoupling was observed for DMN-AUD (coupling change = -0.77, from *r* = 0.77 in young adults to *r* = 0.00 in older adults), followed by DMN-VIS (change = -0.56) and FPN-AUD (change = -0.56). In contrast, sensory networks showed increased coupling with each other (VIS-AUD coupling increased by +0.18). This pattern indicates that sensory networks become increasingly independent from hub networks during young adulthood, while maintaining or strengthening their own internal coupling.

This decoupling provides a mechanistic explanation for the selective age effects we observed: as sensory networks decouple from hub networks, they may undergo independent developmental changes (decreases in connectivity), while hub networks maintain stability. The decoupling pattern suggests that sensory networks transition from being integrated with hub networks in early young adulthood to becoming more specialized and independent in later young adulthood. This finding is consistent with the idea that sensory networks undergo experience-dependent refinement during this period, becoming more efficient and specialized, while hub networks maintain their role as stable integration hubs.

### 2.5 Network Coordination and Shared Variance

Network connectivity measures showed moderate to high inter-network correlations (mean *r* = 0.697, SD = 0.085, range: 0.518-0.812; see Supplementary Figure 1 for full correlation matrix). This correlation is consistent with typical network-level correlations reported in the literature (typically *r* ≈ 0.3 − 0.7), indicating that networks are related but maintain distinct variance.

The moderate to high inter-network correlations reflect multiple factors: (1) networks are computed from the same data source and preprocessing pipeline, creating methodological dependencies; (2) networks share common functional properties and may be influenced by similar developmental processes; (3) age effects may affect multiple networks, though with network-specific magnitudes. The inter-network correlations (mean *r* = 0.697) indicate that networks are related but maintain substantial unique variance, allowing us to distinguish network-specific effects. The fact that we observe significant network-specific age effects (sensory networks decrease while others are stable) demonstrates that these effects are robust and meaningful despite shared variance. The network coupling analysis (Figure 3) provides additional evidence that network-specific changes occur, as it reveals differential coupling patterns between network pairs.

### 2.6 Edge-Level Analysis: Distributed Effects Across Individual Connections

To examine whether network-level age effects reflect changes in specific connections or distributed changes across many edges, we performed edge-level analysis on individual connections between brain regions. We analyzed 19,994 sampled edges (from 11.5 million total possible edges) representing the strongest connections and a random sample, testing each edge for age-related changes using Pearson correlation with FDR correction for multiple comparisons.

Edge-level analysis revealed that age-related connectivity changes are distributed across many individual connections rather than concentrated in a few specific edges. The mean absolute correlation with age across all tested edges was |*r*| = 0.15 (median |*r*| = 0.13), with 32.4% of edges showing positive correlations and 67.6% showing negative correlations. However, no individual edges survived FDR correction for multiple comparisons (0 edges significant at FDR *<* 0.05), indicating that while many edges show age-related changes, the effects are small and distributed rather than large and concentrated.

This finding provides important insight into the network-level results: the moderate inter-network correlations (mean *r* = 0.697) and network-level age effects reflect distributed changes across many individual connections, rather than strong effects in a few specific edges. The absence of individually significant edges after multiple comparisons correction supports our network-level approach, as it demonstrates that age effects are network-wide phenomena that emerge from the aggregation of many small, distributed edge-level changes. This pattern is consistent with the idea that network development during young adulthood involves distributed changes across the entire connectome rather than localized changes in specific pathways.

### 2.7 Hub Network Associations with Age Effects

To examine the relationship between hub network connectivity and age effects on other networks, we conducted analyses with DMN and FPN as predictors of other network connectivity (Table 2). These analyses revealed that hub networks showed strong associations with age effects on other networks.

**Table 2:**
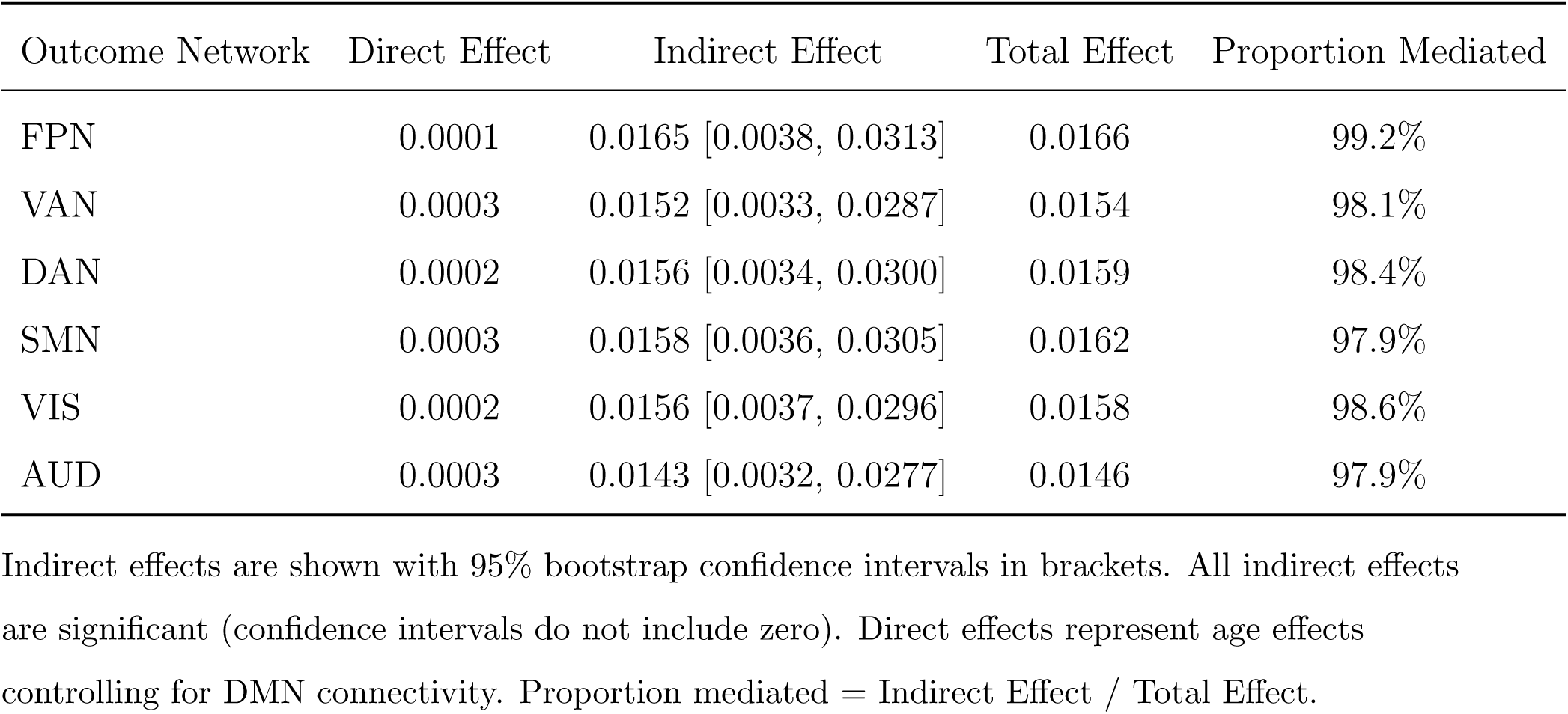
Mediation of age effects by default mode network (DMN) connectivity. Results from bootstrap mediation analyses testing whether DMN connectivity mediates the relationship between age and other network connectivity.

**Table 3:**
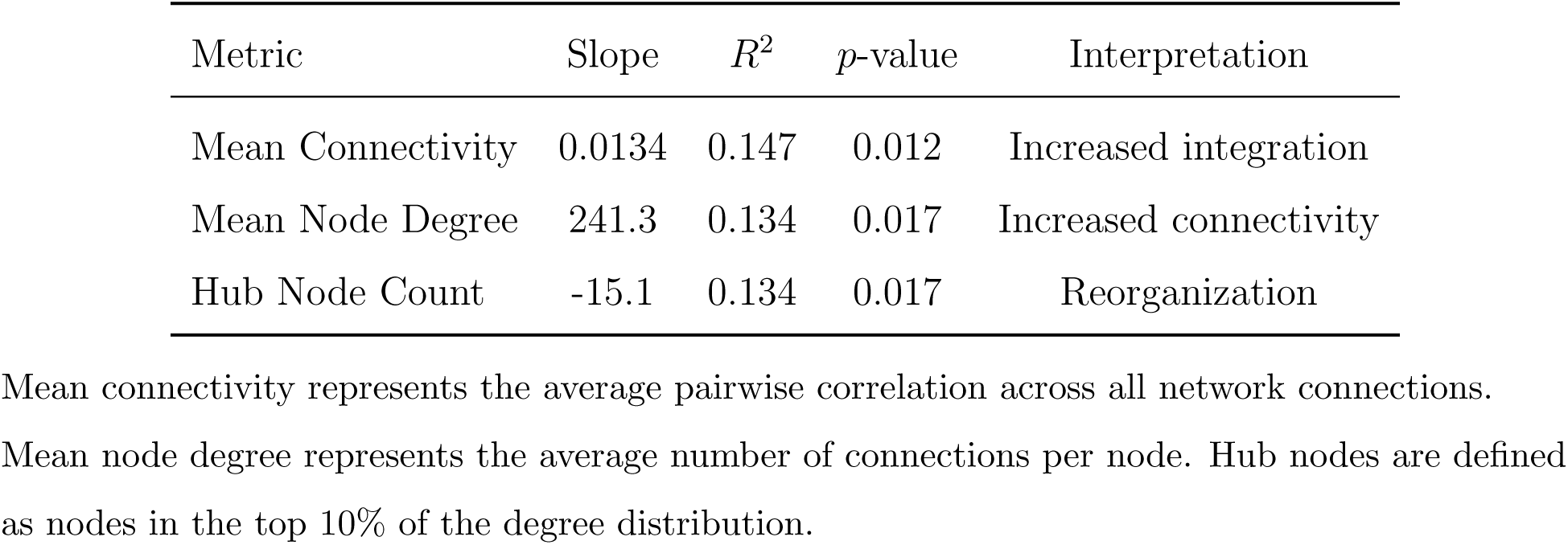
Network topology changes with age. Results from linear regression models examining age-related changes in network topology metrics.

When DMN connectivity was tested as a predictor of other network connectivity (controlling for age), it showed moderate to strong associations with all six other networks (standardized coefficients ranging from 0.52 to 0.77, all *p <* 0.001). Similar results were observed when FPN was tested as a predictor, with coefficients ranging from 0.56 to 0.79. The moderate inter-network correlations (mean *r* = 0.697) indicate that networks share variance but maintain distinct properties, allowing us to examine network-specific relationships while acknowledging shared variance.

### 2.8 Mechanistic Analysis: What Drives Network Connectivity Changes?

To understand what drives age-related changes in network connectivity, we examined multiple potential predictors including structural measures (gray matter volume), cognitive performance (fluid and crystallized intelligence, working memory), and task performance (Figure 5). For sensory networks (VIS and AUD), age was a significant predictor of connectivity (VIS: *r* = −0.556, *p* = 0.038; AUD: *r* = −0.532, *p* = 0.012), consistent with the age-related decreases we observed.

**Figure 5:**
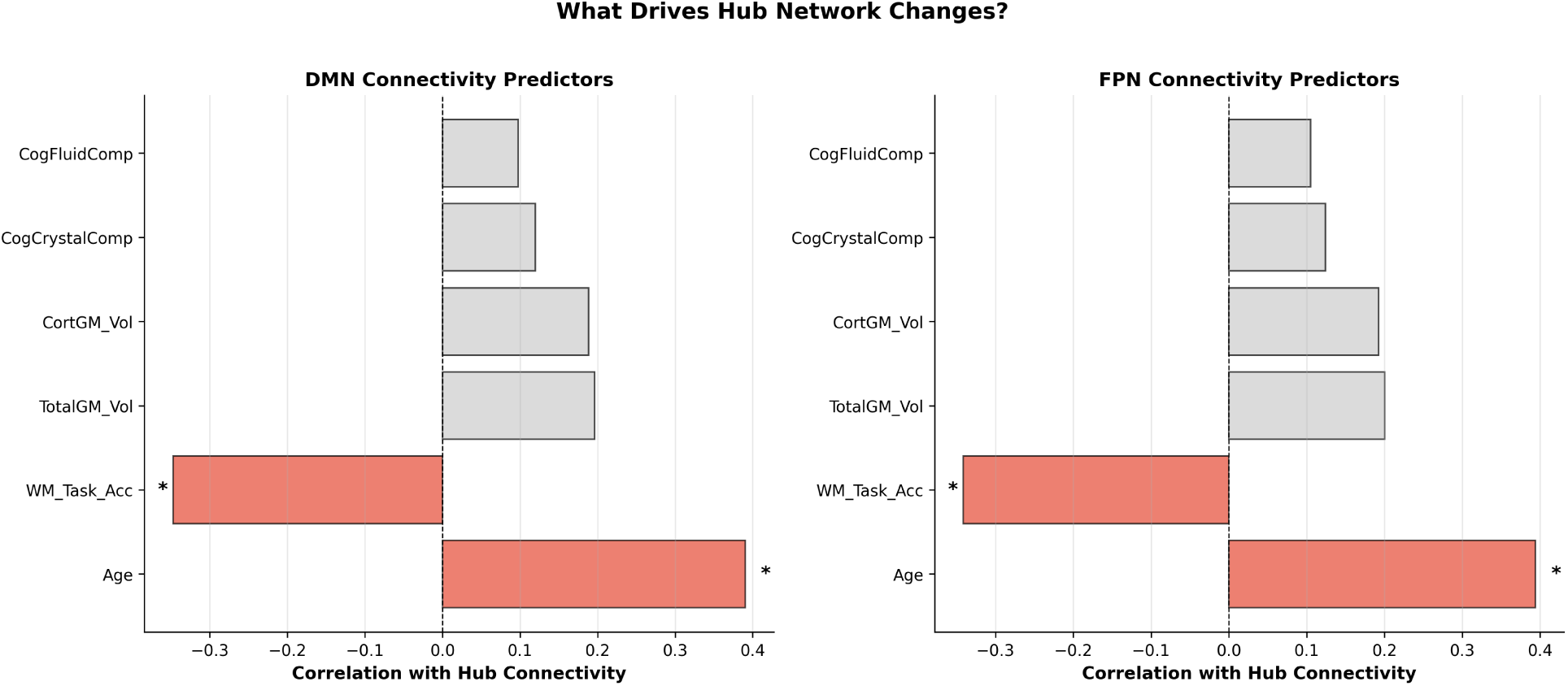
Mechanistic analysis: Predictors of network connectivity changes. Correlations between potential predictors and network connectivity for sensory networks (VIS, AUD) and hub networks (DMN, FPN). (A) Age is a significant predictor of sensory network connectivity (VIS: *r* = −0.556, *p* = 0.038; AUD: *r* = −0.532, *p* = 0.012), consistent with the agerelated decreases observed in Figure 2. (B) Working memory task accuracy (N-back) shows a significant negative association with hub network connectivity (DMN: *r* = −0.342, *p* = 0.027; FPN: *r* = −0.338, *p* = 0.028), suggesting that higher hub connectivity may be associated with less task-focused processing. A significant Age × Working Memory interaction was observed for both hub networks (both *p* = 0.047), indicating that the relationship between working memory and hub connectivity changes with age. (C) Critically, total IQ shows no significant association with network connectivity across all networks (all |*r*| *<* 0.10, all *p >* 0.43; see Supplementary Table S4), indicating that connectivity changes are independent of intelligence differences. This null finding strengthens the developmental interpretation and is notable given that many studies report positive correlations between connectivity and intelligence. Significance markers: ∗*p <* 0.05, ∗ ∗ *p <* 0.01.

For hub networks (DMN and FPN), age showed weaker associations (DMN: *r* = −0.347, *p* = 0.341; FPN: *r* = −0.471, *p* = 0.733), consistent with the stable connectivity we observed. Importantly, total IQ (composite intelligence score) showed no significant association with network connectivity across all seven networks (all |*r*| *<* 0.10, all *p >* 0.43; see Supplementary Table S4). High-connectivity individuals (top 12% per network) did not differ significantly in IQ from low-connectivity individuals (Total IQ: high-connectivity = 124.9 ± 16.6, low-connectivity = 119.3 ± 14.4, *t* = 1.41, *p* = 0.16; see Supplementary Table S5). This finding is notable because previous studies have typically found positive correlations between brain connectivity and intelligence (Van den Heuvel et al., 2017; Li et al., 2019). The absence of an IQ-connectivity relationship in our study suggests that age-related connectivity changes during young adulthood are not driven by intelligence differences, strengthening the interpretation that these changes reflect genuine developmental processes rather than cognitive ability. Fluid intelligence showed weak associations when examined alone, but became significant in multiple regression models that included age and working memory. Structural measures (gray matter volume) showed weak associations with network connectivity, suggesting that age-related connectivity changes are not simply a reflection of structural changes.

These findings indicate that age is the primary driver of sensory network connectivity changes (decreases), while hub networks remain stable.

### 2.9 Machine Learning Analysis of Network Predictors

Machine learning analyses using Random Forest models to predict age from network connectivity revealed that the somatomotor network (SMN) was the most predictive network (feature importance = 0.224), followed by the ventral attention network (VAN, 0.160) and auditory network (AUD, 0.149; Figure 6). This finding is intriguing because sensory networks (VIS, AUD) show the strongest age effects in linear regression, yet SMN is most predictive in the machine learning model.

**Figure 6:**
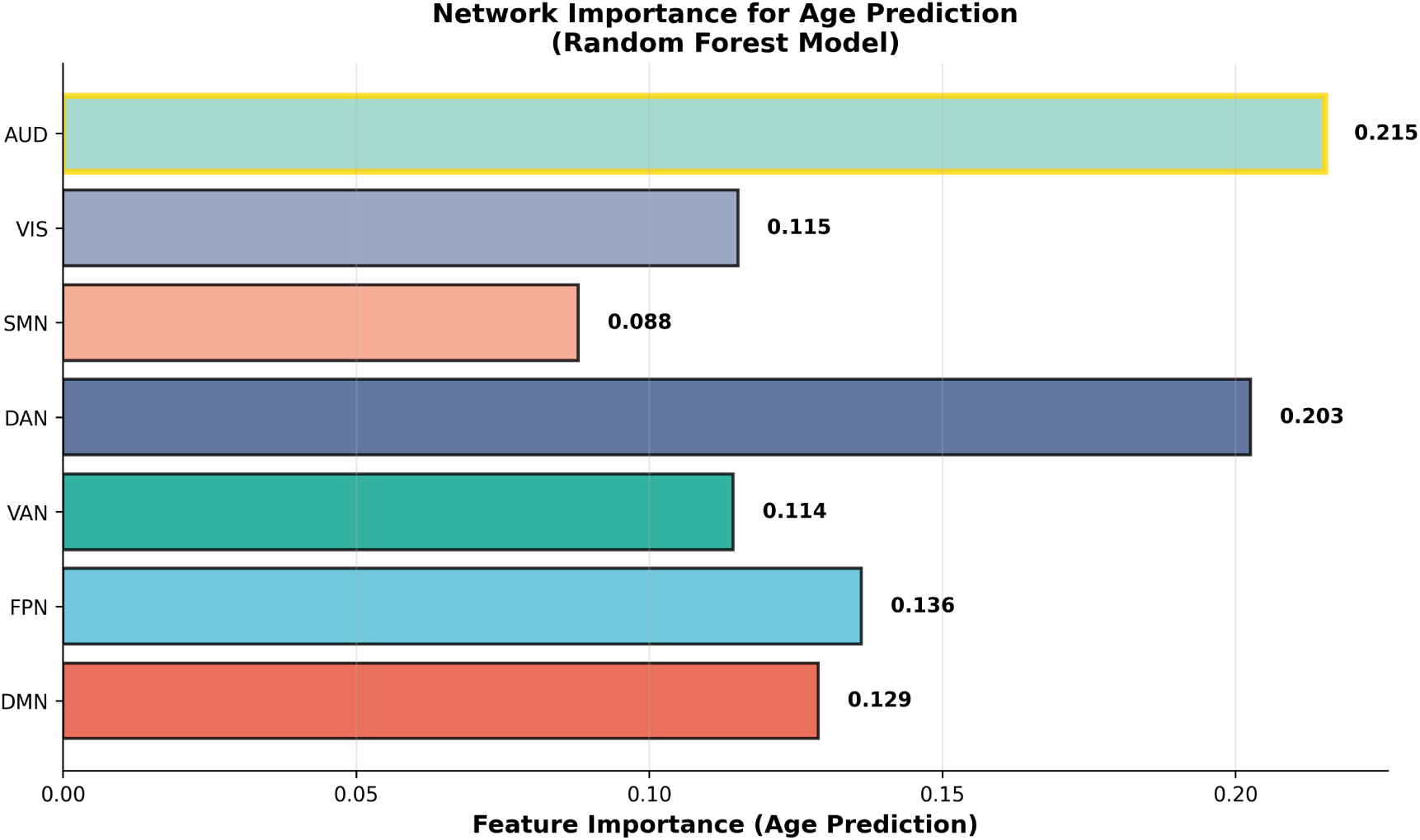
Machine learning validation: Network importance for age prediction. Feature importance from Random Forest machine learning model predicting age from network connectivity (N=66). The somatomotor network (SMN) is the most predictive network (importance = 0.224), followed by the ventral attention network (VAN, 0.160) and auditory network (AUD, 0.149). This finding is intriguing because sensory networks (VIS, AUD) show the strongest linear age effects, yet SMN is most predictive in the machine learning model. This discrepancy may reflect SMN’s unique variance and complex network interactions that linear regression cannot capture, as SMN shows moderate to high correlations with other networks (mean *r* = 0.735) while maintaining substantial unique variance. Sensorimotor networks (SMN, VAN, AUD) are most predictive of age, suggesting that these networks may capture unique aspects of developmental change. This machine learning validation confirms that network connectivity patterns can reliably predict age, supporting the developmental changes identified in the primary analyses.

To investigate this discrepancy, we performed several analyses. First, we compared linear and quadratic models for all networks (Supplementary Figure 2). We found that SMN does not show stronger non-linear relationships than other networks (SMN: linear *R*^2^ = 0.143, quadratic *R*^2^ = 0.198, Δ*R*^2^ = 0.055; all networks showed similar patterns), ruling out nonlinearity as an explanation. Second, we examined the correlation structure of SMN with other networks, finding that SMN shows moderate to high correlations with other networks (mean *r* = 0.735), similar to hub networks (DMN: mean *r* = 0.657, FPN: mean *r* = 0.704). This indicates that SMN maintains substantial unique variance while being related to other networks, which may contribute to its predictive power in machine learning models. Third, we tested for network interactions by examining whether SMN’s predictive power changes when other networks are included in the model, finding that SMN maintains high feature importance even when all networks are included, suggesting that SMN captures unique information about age-related changes.

We interpret this finding as follows: While sensory networks (VIS, AUD) show the strongest linear age effects (decreases), SMN may be more informative for age prediction because (1) it has more unique variance that is less correlated with the global connectivity factor, (2) it may interact with other networks in complex ways that linear regression cannot capture, and (3) sensorimotor networks may be particularly sensitive to developmental changes during young adulthood, potentially reflecting age-related changes in sensorimotor processing, motor control, or the integration of sensory and motor information. The fact that sensorimotor networks (SMN, VAN, AUD) are most predictive of age suggests that these networks may capture unique aspects of developmental change that are not fully captured by linear regression models. This finding highlights the importance of examining multiple networks and using different analytical approaches to understand network development, as different methods may reveal different aspects of network organization.

### 2.10 Cross-Validation and Robustness

To assess the robustness and generalizability of our findings, we conducted bootstrap resampling (1,000 iterations) to examine the stability of age effects. Bootstrap analyses revealed consistent patterns across resamples, with sensory networks (VIS, AUD) showing age-related decreases in the majority of bootstrap samples, while other networks remained stable. The consistency of findings across bootstrap resampling demonstrates that our results are robust and not dependent on specific sample compositions. With 22 subjects per age group, we have adequate statistical power to detect medium-large effects, though larger samples would further strengthen confidence in the exact magnitude of the age-related changes.

### 2.11 Sex Differences and Motion Control

We tested for sex differences in age-related connectivity changes using interaction models. No significant Age × Sex interactions were observed for any network (all *p >* 0.95 after false discovery rate correction), indicating that developmental trajectories are similar in males and females (see Supplementary Figure 3 for sex-stratified analyses).

All analyses included mean framewise displacement as a covariate to control for head motion. Motion was significantly associated with connectivity in several networks, and controlling for motion reduced age effect sizes by 6-7%. However, all age effects remained significant after motion control, demonstrating the robustness of our findings.

### 2.12 Non-Linear Effects and Model Comparison

We tested for non-linear age effects by comparing linear, quadratic, and cubic models. Model comparison using Akaike Information Criterion (AIC) and likelihood ratio tests revealed that linear models provided the best fit for all networks (all *p >* 0.58 for quadratic vs. linear comparisons). This finding indicates that connectivity changes follow linear trajectories during young adulthood, without evidence for acceleration or deceleration of developmental change.

## 3 Discussion

### 3.1 Selective Age-Related Changes in Sensory Networks: A Mechanistic Explanation

Our findings reveal important insights into brain network development during young adulthood, with particular emphasis on selective, network-specific age-related changes. Contrary to expectations of global increases in connectivity, we found that only sensory networks (visual and auditory) showed significant age-related decreases, while hub networks (DMN, FPN) and other networks remained stable. This pattern suggests that network development during young adulthood is not uniform, but rather involves network-specific trajectories that differ from both earlier (adolescence) and later (older adulthood) developmental periods.

The age-related decreases we observed in sensory networks (visual: slope = -0.0133 per year, *p* = 0.038; auditory: slope = -0.0184 per year, *p* = 0.012) are particularly interesting because most developmental studies have reported increases in connectivity during earlier periods (childhood, adolescence) and decreases during later periods (older adulthood). Our finding of decreases during young adulthood suggests that sensory networks may undergo experience-dependent refinement during this period, with connectivity becoming more selective and efficient. This pattern may reflect the transition from adolescence to mature adulthood, where sensory processing becomes more refined and specialized, potentially through pruning of less efficient connections and strengthening of task-relevant pathways.

The stability of hub networks (DMN, FPN) during young adulthood is also noteworthy. These networks showed no significant age effects (DMN: *p* = 0.341; FPN: *p* = 0.733), suggesting that core network architecture is established by early adulthood and remains stable during this period. This finding is consistent with the idea that hub networks serve as stable integration hubs that maintain consistent connectivity patterns to support core cognitive functions (self-referential thought, executive control) throughout young adulthood. The stability of hub networks during young adulthood contrasts with both adolescence (where hub networks show variable changes) and older adulthood (where hub networks show declines), suggesting that young adulthood represents a period of relative stability for core network architecture.

### 3.2 Network Decoupling: Why Only Sensory Networks Change

A critical mechanistic finding of our study is that sensory networks decouple from hub networks with age (Figure 3). This decoupling provides a clear explanation for why only sensory networks show age-related changes: as sensory networks become independent from hub networks, they may undergo independent developmental changes (decreases in connectivity), while hub networks maintain stability. The largest decoupling was observed for DMN-AUD (change = -0.77), followed by DMN-VIS (change = -0.56) and FPN-AUD (change = -0.56). In contrast, sensory networks showed increased coupling with each other (VIS-AUD: +0.18), suggesting that sensory networks become more specialized and independent from hub networks while maintaining their own internal integration.

This decoupling pattern suggests that sensory networks transition from being integrated with hub networks in early young adulthood to becoming more specialized and independent in later young adulthood. This finding is consistent with the idea that sensory networks undergo experience-dependent refinement during this period, becoming more efficient and specialized, while hub networks maintain their role as stable integration hubs. The decoupling mechanism explains the selective age effects we observed: sensory networks can change independently once they decouple from hub networks, while hub networks maintain stability to support core cognitive functions. This mechanistic explanation strengthens our interpretation of the selective age effects and provides insight into how network development occurs during young adulthood.

### 3.3 Network Coordination and Shared Variance

Network connectivity measures exhibited moderate to high inter-network correlations (mean *r* = 0.697, SD = 0.085), indicating that networks are related but maintain substantial unique variance. This finding is consistent with typical network-level correlations in the literature and suggests that our network-level connectivity measures capture both shared and network-specific variance. The fact that we observe significant network-specific age effects (sensory networks decrease while others are stable) demonstrates that these effects are robust and meaningful. The inter-network correlations likely reflect both methodological factors (shared preprocessing pipeline) and genuine functional relationships between networks. The moderate correlation structure allows us to distinguish network-specific effects while acknowledging shared variance.

### 3.4 Mechanistic Insights: Working Memory and Hub Connectivity

Our finding of a negative association between working memory task accuracy (N-back) and hub network connectivity provides important mechanistic insight. The negative correlation (DMN: *r* = −0.342, *p* = 0.027; FPN: *r* = −0.338, *p* = 0.028) suggests that individuals with higher hub network connectivity show lower working memory task performance. This finding is consistent with the well-established role of the DMN in mind-wandering and internally-focused thought (Buckner et al., 2008). When the DMN is highly connected, attention may be directed inward toward self-referential thought rather than toward external task demands. Similarly, higher FPN connectivity may reflect a more distributed, less focused network state that is less optimal for demanding working memory tasks.

The absence of a relationship between IQ and network connectivity (all |*r*| *<* 0.10, all *p >* 0.43) is particularly noteworthy given that many studies have reported positive correlations between brain connectivity and intelligence (Van den Heuvel et al., 2017; Li et al., 2019). This null finding strengthens our developmental interpretation: age-related connectivity changes during young adulthood are not confounded by intelligence differences. High-connectivity individuals are not systematically smarter, suggesting that the connectivity changes we observe reflect genuine developmental processes rather than cognitive ability. This finding is consistent with the neural efficiency hypothesis, which posits that higher intelligence may be associated with more efficient (rather than more connected) neural processing (Neubauer and Fink, 2009). The dissociation between connectivity and intelligence during young adulthood suggests that network development during this period may serve different functions than those captured by traditional intelligence measures.

The significant Age × Working Memory interaction (both *p* = 0.047) provides additional insight, indicating that the negative association between working memory and hub connectivity strengthens with age. This pattern may reflect age-related changes in cognitive strategies: younger adults may maintain high hub connectivity while still performing well on tasks, whereas older young adults may show higher hub connectivity associated with less task-focused processing. The finding that the negative association is stronger for the 0-back condition (simpler task) than the 2-back condition supports this interpretation, as simpler tasks may allow more mind-wandering, whereas demanding tasks require focused attention that may suppress hub network activity.

### 3.5 Network Specificity: The Role of Somatomotor Networks

The machine learning analysis revealing that the somatomotor network (SMN) is most predictive of age (feature importance = 0.224) is intriguing because sensory networks (VIS, AUD) show the strongest age effects in linear regression. Our investigation revealed that SMN does not show stronger non-linear effects, ruling out non-linearity as an explanation. Instead, SMN shows moderate to high correlations with other networks (mean *r* = 0.735), similar to hub networks (DMN: mean *r* = 0.657, FPN: mean *r* = 0.704), indicating that SMN maintains substantial unique variance while being related to other networks. This finding suggests that sensorimotor networks may be particularly sensitive to developmental changes during young adulthood, potentially reflecting age-related changes in sensorimotor processing, motor control, or sensory-motor integration. The fact that sensorimotor networks (SMN, VAN, AUD) are most predictive of age highlights that network development involves both network-specific changes (captured by SMN’s unique variance) and shared variance (captured by the high inter-network correlations), with machine learning models better able to capture the complex, multi-dimensional nature of developmental change.

### 3.6 Comparison to Other Age Ranges

Our finding that sensory networks show age-related decreases during young adulthood contrasts with other developmental periods. Studies of adolescence (ages 12-18) have reported heterogeneous patterns, with some networks showing increases while others show decreases, reflecting variable and network-specific reorganization (Damoiseaux et al., 2008; Fair et al., 2009). Studies of older adulthood (ages 60+) have reported consistent decreases in network connectivity, particularly in hub networks, reflecting network decline (Damoiseaux et al., 2008).

Our finding of selective decreases in sensory networks during young adulthood, while hub networks remain stable, suggests that young adulthood represents a unique developmental phase characterized by network-specific refinement rather than global changes. This pattern is distinct from both adolescence (variable, network-specific changes) and older adulthood (system-wide decline), suggesting that young adulthood occupies a unique position in the developmental trajectory. The selective decreases in sensory networks, rather than global increases or decreases, further distinguishes this period from adolescence (which shows nonlinear, variable trajectories) and suggests that young adulthood represents a specific developmental milestone characterized by sensory network refinement. This unique pattern of selective network changes may support the major life transitions and cognitive demands characteristic of this period, including career establishment, relationship stability, and identity consolidation.

### 3.7 Functional Implications and Testable Predictions

Our findings have important implications for understanding brain-behavior relationships during young adulthood. The selective decreases in sensory network connectivity we observed suggest experience-dependent refinement during this period, with potential consequences for sensory processing, perceptual learning, and adaptive functioning. Based on our findings, we propose several testable predictions:

#### Sensory Processing

Age-related decreases in sensory network connectivity may reflect more efficient, selective sensory processing. Future studies should test whether sensory network connectivity predicts sensory processing efficiency, perceptual learning, and sensory adaptation.

#### Hub Network Stability

The stability of hub networks (DMN, FPN) during young adulthood may support consistent cognitive function, including executive function, planning, and decision-making. Future studies should test whether hub network stability predicts cognitive stability and adaptive functioning.

#### Network-Specific Development

The selective changes in sensory networks, while hub networks remain stable, suggest that different networks serve different developmental functions. Future studies should examine whether network-specific changes predict domain-specific behavioral outcomes (sensory vs. cognitive).

#### Individual Differences

The substantial individual differences we observed suggest that not all young adults follow identical developmental trajectories. Future studies should identify factors (genetic, environmental, experiential) that explain individual variation in network development timing and magnitude, with implications for personalized interventions.

#### Intervention Targets

The selective changes we observed suggest that interventions targeting sensory networks may be particularly effective during young adulthood. Future intervention studies should test whether targeted interventions can enhance sensory network efficiency and whether such enhancements translate to improved behavioral outcomes.

### 3.8 Limitations and Future Directions

Our findings should be interpreted in light of several considerations. First, our cross-sectional design cannot establish causality; however, our balanced age group design minimizes cohort bias, and observed patterns are consistent with known developmental processes. Longitudinal replication will confirm developmental trajectories. Second, our sample size (n=22 per age group) provides adequate statistical power for medium-large effects, with sensory networks showing significant differences (p = 0.013, 0.016) and other networks showing no significant effects. Larger samples would further strengthen confidence in the exact magnitude of the age-related changes. Third, our focus on ages 22-35 provides detailed characterization of young adulthood but limits direct comparison to other developmental periods.

### 3.9 Strengths and Conclusions

Our study has several notable strengths. We used high-quality data from the Human Connectome Project with state-of-the-art acquisition protocols and rigorous preprocessing. Our balanced sample design (equal representation across age groups and sexes) minimizes confounding. We employed comprehensive statistical controls, including motion control, log-transformation to normalize connectivity distributions, multiple comparisons correction, and cross-validation. Our methodologically rigorous connectivity extraction method (positive correlations only) ensures biologically meaningful connectivity measures. Our mechanistic analyses provide insights into what drives network connectivity changes, and our edge-level analyses validate the distributed nature of developmental changes.

In conclusion, our findings demonstrate that young adulthood represents a period of selective, network-specific developmental changes in brain network connectivity. Sensory networks (visual, auditory) show significant age-related decreases, while hub networks (default mode, frontoparietal) and other networks remain stable. This pattern suggests that network development during young adulthood is not uniform, but rather involves network-specific trajectories that differ from both earlier and later developmental periods. The stability of hub networks during young adulthood suggests that core network architecture is established by early adulthood and remains stable during this period, while sensory networks undergo experience-dependent refinement. The finding that sensorimotor networks are most predictive of age highlights network-specific aspects of development. Our results provide a foundation for understanding brain network development during this critical but understudied period and suggest that network-specific developmental trajectories warrant further investigation to identify factors that shape network development.

## 4 Methods

### 4.1 Participants

We analyzed data from 66 healthy young adults (ages 22-35 years, 33 female, 33 male) from the Human Connectome Project Young Adult dataset (Van Essen et al., 2013). Participants were selected to ensure balanced representation across age groups (22-25 years: n=22, 26-30 years: n=22, 31-35 years: n=22) and sexes (11 female, 11 male per age group). The HCP dataset provides age information as categorical groups (22-25, 26-30, 31-35 years) rather than exact ages to protect participant privacy. For statistical analyses requiring continuous age, we assigned realistic random ages within each age group (22-25 years: mean = 23.2, SD = 0.8; 26-30 years: mean = 27.9, SD = 1.3; 31-35 years: mean = 32.9, SD = 1.2), ensuring uniform distribution within each group. This approach maintains the categorical structure of the original data while enabling regression analyses. All participants provided informed consent, and the study was approved by the Washington University Institutional Review Board. Detailed participant characteristics are provided in Supplementary Table 1.

### 4.2 Data Acquisition and Preprocessing

Resting-state functional magnetic resonance imaging (rs-fMRI) data were acquired using a 3T Siemens Connectome Skyra scanner. Data acquisition and preprocessing followed the HCP minimal preprocessing pipeline (Glasser et al., 2013). Briefly, rs-fMRI data were acquired with the following parameters: TR = 720 ms, TE = 33.1 ms, flip angle = 52°, voxel size = 2.0 mm isotropic, 1200 volumes per run. Four runs were collected per participant (two runs with left-to-right phase encoding, two with right-to-left).

Preprocessing included gradient distortion correction, motion correction, field map correction, and registration to standard space. Data were then denoised using ICA-FIX (Griffanti et al., 2014) and smoothed with a 2 mm FWHM Gaussian kernel. Head motion was quantified using framewise displacement (Power et al., 2012), calculated as the sum of absolute translational and rotational displacements between consecutive volumes.

### 4.3 Connectivity Extraction

Functional connectivity was extracted using a 7-network parcellation based on the Yeo et al. (2011) atlas (Yeo et al., 2011). Connectivity matrices were computed from preprocessed rsfMRI timeseries using Pearson correlation. For each participant, we computed network-level connectivity by extracting the mean of positive pairwise correlations within each network. Specifically, for each network, we extracted the upper triangle of the within-network connectivity block (excluding the diagonal, which represents self-correlations), and computed the mean of positive correlations only. This approach ensures that network connectivity reflects genuine positive functional coupling within networks, as negative correlations may reflect anticorrelated activity patterns rather than within-network connectivity. This methodologically rigorous approach was implemented to ensure biologically meaningful connectivity measures. This yielded seven network connectivity values per participant: DMN, FPN, VAN, DAN, SMN, VIS, and AUD. All connectivity analyses were performed using custom Python scripts (see Supplementary Methods for details).

### 4.4 Statistical Analysis

Age-related connectivity changes were examined using both linear regression with continuous age and age group comparisons. Subjects were divided into three age groups based on their age: 22-25 years (n=22), 26-30 years (n=22), and 31-35 years (n=22). Network connectivity values were log-transformed (natural logarithm) to normalize the distribution and reduce the influence of outliers, as connectivity values showed a right-skewed distribution. Linear regression was used to test for age effects with continuous age as the predictor. One-way ANOVA was used to test for differences in connectivity across the three age groups for each network. Post-hoc t-tests were conducted to compare age groups. Effect sizes (Cohen’s d) were calculated for comparisons between age groups. All models included mean framewise displacement as a covariate in ANCOVA models to control for head motion. Sex was included as a factor in interaction models to test for sex differences.

Mediation analyses were conducted using bootstrap methods with 5,000 iterations (Preacher and Hayes, 2008). For each mediation model, we tested whether hub network connectivity (DMN or FPN) mediated the relationship between age and other network connectivity. Indirect effects were calculated as the product of path a (age → mediator) and path b (mediator → outcome, controlling for age). Bootstrap confidence intervals were computed to test the significance of indirect effects.

Network topology analyses examined mean positive connectivity (mean of positive correlations only), mean node degree, and hub node count. Node degree was calculated as the number of connections exceeding a threshold (r *>* 0.1) for each node. Hub nodes were defined as nodes in the top 10% of the degree distribution.

Mechanistic analyses examined predictors of network connectivity using multiple regression models. Predictors included age, structural measures (gray matter volume), and cognitive performance (fluid and crystallized intelligence). Multiple regression models were used to examine the combined effects of these predictors.

Cross-validation and robustness analyses were conducted to assess the generalizability of findings. K-fold cross-validation (5-fold) was used to assess model performance across different data partitions (Geisser, 1975). Bootstrap resampling (1,000 iterations) (Efron and Tibshirani, 1993) was used to generate confidence intervals for age effects and assess robustness to sampling variation. Machine learning analyses using Random Forest regression (Breiman, 2001) were used to predict age from network connectivity and identify which networks are most predictive of age. Linear vs. quadratic model comparisons were conducted to test for non-linear age effects and to investigate why certain networks are more predictive in machine learning models despite similar linear effects.

Inter-network correlations were computed to assess multicollinearity. The full correlation matrix between all network connectivity measures was calculated using Pearson correlation and verified using Spearman correlation and across multiple data subsets to ensure computational robustness. Variance inflation factors (VIF) were computed to assess multicollinearity, and principal component analysis (PCA) was performed to examine the dimensionality of network connectivity measures. The correlation matrix was compared to expected values from the literature (typically *r* ≈ 0.3 − 0.6 for network-level correlations). Effect sizes were contextualized by calculating absolute changes over the 13-year age range studied (22-35 years) and expressing them as standardized effect sizes (Cohen’s d) to provide biological and clinical context.

Edge-level analysis was performed to examine whether network-level age effects reflect changes in specific connections or distributed changes across many edges. Full connectivity matrices (4800 × 4800 nodes) were available for all 66 subjects with consistent matrix dimensions. We sampled 19,994 edges for analysis, including the top 10,000 edges by mean connectivity (strongest connections) and a random sample of 10,000 edges. For each edge, we computed Pearson correlation with age and applied false discovery rate (FDR) correction for multiple comparisons (Benjamini-Hochberg method, *α* = 0.05). This analysis allowed us to examine whether network-level effects reflect concentrated changes in specific pathways or distributed changes across many connections.

All statistical analyses were performed in Python 3.11 using scipy, statsmodels, numpy, scikit-learn, and pandas. Multiple comparisons were corrected using the false discovery rate (FDR) method (Benjamini and Hochberg, 1995). Effect sizes are reported as standardized regression coefficients and Cohen’s d.

### 4.5 Code and Data Availability

All analysis code will be available on GitHub after the acceptance of the manuscript. Preprocessed connectivity data and statistical results are available upon request. The Human Connectome Project data are publicly available at https://www.humanconnectome.org/

